# Thyroid hormone and thyromimetics inhibit myelin and axonal degeneration and oligodendrocyte loss in EAE

**DOI:** 10.1101/2020.12.20.423638

**Authors:** P Chaudhary, GH Marracci, E Calkins, E Pocius, AL Bensen, TS Scanlan, B Emery, DN Bourdette

**Author notes:** These authors contributed equally to manuscript (P Chaudhary, DN Bourdette). Corresponding author: Priya Chaudhary, PhD.

## Abstract

We have previously demonstrated that thyromimetics stimulate oligodendrocyte precursor cell differentiation and promote remyelination in murine demyelination models. We investigated whether a thyroid receptor-beta selective thyromimetic, sobetirome (Sob), and its CNS-targeted prodrug, Sob-AM2, could prevent myelin and axonal degeneration in experimental autoimmune encephalomyelitis (EAE). Compared to controls, EAE mice receiving triiodothyronine (T3, 0.4mg/kg), Sob (5mg/kg) or Sob-AM2 (5mg/kg) had reduced clinical disease and, within the spinal cord, less tissue damage, more normally myelinated axons, fewer degenerating axons and more oligodendrocytes. T3 and Sob also protected cultured oligodendrocytes against cell death. Thyromimetics thus might protect against oligodendrocyte death, demyelination and axonal degeneration as well as stimulate remyelination in multiple sclerosis.

**Highlights:**

- Thyroid hormone, the thyromimetic Sob and its CNS penetrating prodrug, Sob-AM2, reduce disease severity, reduce myelin and axonal degeneration and protect oligodendrocytes in EAE.
- The benefits of Sob and Sob-AM2 may be via direct protective effects on oligodendrocytes and reduction in activity of microglia/macrophages.

## 1. Introduction

Thyroid hormone (TH) regulates oligodendrocyte differentiation and myelination during development and stimulates remyelination (Billion *et al.* 2002; Schoonover *et al.* 2004; Calza *et al.* 2018; Hartley *et al.* 2019). The active form of TH, 3,5,3’-triiodothyronine (T3), binds and activates thyroid hormone receptors TRα and TRβ that act as transcription factors to influence gene expression of thyroid hormone regulated genes (Bernal 2007). Importantly, TH accelerates remyelination in the cuprizone and lysolecithin models of demyelination (Zendedel *et al.* 2016; Hartley *et al.* 2019). TH treatment also provides beneficial effects in experimental autoimmune encephalomyelitis (EAE), displaying anti-inflammatory effects and reduction in demyelination and axonal injury (Calza *et al.* 2002; Dell’Acqua *et al.* 2012). Despite its therapeutic benefit in models of multiple sclerosis (MS), TH cannot be used therapeutically in MS because there is likely no margin separating the beneficial CNS effects from systemic thyrotoxicosis (Wooliscroft *et al.* 2020).

The TRβ selective thyroid hormone agonist sobetirome (Sob) (also called GC-1) lacks the systemic toxic effects of TH but retains its therapeutic benefits (Scanlan 2010). To enhance the CNS effects of Sob, we previously developed and evaluated CNS-penetrating prodrugs, including Sob-AM2, that cross the blood brain barrier and within the CNS are hydrolyzed to produce high levels of Sob (Meinig *et al.* 2017, 2018; Ferrara *et al*. 2017; Bárez-López *et al*. 2018; Ferrara and Scanlan 2020). Compared to Sob, an equivalent bioavailable fraction of Sob-AM2 delivers more than ten times the thyromimetic exposure to the CNS while masking thyromimetic activity in the periphery, thus increasing therapeutic index (Meinig *et al.* 2018). We previously showed that Sob and Sob-AM2 stimulated oligodendrocyte precursor cell (OPC) differentiation and remyelination in a novel genetic mouse model of demyelination and Sob promoted remyelination in the cuprizone and lysolecithin models (Hartley *et al.* 2019). In the present study, we assessed for the first time the therapeutic benefits of Sob and Sob-AM2 in an EAE model of MS.

## 2. Methods

### 2.1 EAE induction and drug delivery

The VA Portland Health Care System (VAPORHCS) IACUC committee approved all experiments. C57BL/6 female mice (The Jackson Laboratory, Bar Harbor, ME, ages 8-10 weeks) were immunized with 200μg of myelin oligodendrocyte glycoprotein (MOG) 35-55 peptide (PolyPeptide Laboratories, San Diego, CA) in complete Freund’s adjuvant containing 400μg of *Mycobacterium tuberculosis* per mouse (subcutaneous injection of 0.2ml volume). Pertussis toxin (List Biological labs Inc, Campbell, CA) was administered via intraperitoneal (ip) injection at day 0 (75ng per mouse) and day 2 (200ng per mouse) after immunization. Mice were scored for clinical signs of EAE daily using a previously described 9-point scale (Forte *et al.* 2007). Scoring was done daily before administration of treatment and without reference to previous EAE scores. Mice received daily ip injections of vehicle (50% DMSO or 8mM NaOH, both in saline) or drug (T3 0.4mg/kg, Sob 5mg/kg, or Sob-AM2 5mg/kg) starting on day 7 before signs of EAE and continued daily treatment through euthanasia on day 21 post-immunization. The experiment was repeated 3 times with 6-8 mice per group for each experiment. After 21 days, mice were euthanized and spinal cords were isolated after perfusion (2% glutaraldehyde and 2% paraformaldehyde, n=2 per group) or immersion fixation (4% paraformaldehyde, n=4-6 per group). Spinal cords were isolated and the tissue was further fixed using microwave fixation (Calkins et al. 2019). Only immersion fixed tissue was used for immunofluorescence. Spinal cords sections from perfused and immersion fixed tissue were used for embedding in plastic which was used for determination of tissue damage and electron microscopy (EM).

### 2.2 Immunofluorescence staining of spinal cord slices

After immersion fixation, lumbar spinal cord was isolated and sectioned. 50μm sections were cut for immunostaining and 350μm sections were used for embedding in plastic. For each spinal cord, two sections (50μm) were randomly chosen and permeabilized with 0.2% Triton X in PBS for 30-45 min and then blocked with 0.5% fish skin gelatin, 3% BSA in PBS for 2 hours at room temperature. The sections were then incubated with primary antibodies: anti CD4 (BD Biosciences, 550280, 1:25 dilution), CD11b (BioRad, Hercules, CA, MCA711, 1:250 dilution), Iba1 (Wako, Richmond, VA, 019-19741, 1:250 dilution), or aspartoacylase (ASPA, Millipore, Burlington, MA, ABN1698, 1:200 dilution) at 4°C overnight. The sections were washed three times and incubated with Alexa fluor secondary antibodies for two hours at room temperature. Donkey anti-rat IgG Alexa Fluor 488, A21208 or Donkey anti-Rabbit IgG Alexa Fluor 555, A-31572 (both from Thermo Fisher, Waltham, MA) were used as secondary antibodies at 1:200 dilution. DAPI (1 μg/ml) was used as a nuclear marker. The sections were mounted in Prolong Gold antifade (Thermo Fisher, P36934) and imaged using Zeiss 780 laser scanning confocal. All sections were processed without knowledge of treatment group.

### 2.3 Light and electron microscopy of plastic embedded spinal cord

Lumbar spinal cord sections (350μm) from immersion fixed and perfused mice were embedded using a previously described protocol (Hartley *et al.* 2019, Calkins *et al.* 2019). Briefly, the spinal cord sections were post-fixed in 1.5% paraformaldehyde, 1.5% glutaraldehyde, 0.05M sucrose, 0.25% calcium chloride in 0.05M sodium cacodylate buffer pH 7.4. All embedding steps were performed using the Biowave Pro + (Ted Pella, Redding, CA). The tissue was processed using 2% osmium-1.5% potassium ferrocyanide, rinsed with distilled water, immersed in 0.5% uranyl acetate, passed through series of acetone solutions (20, 30, 50, 75, 95, 100%) for dehydration, infiltrated with acetone/resin mixtures and finally with resin (Spurr and Eponate 12). The blocks were polymerized in an oven for 1-2 days at 63°C. The blocks were removed from capsules, sectioned semi-thin (0.5μm) for light microscopy and ultra-thin (70nm) for EM. All sections were processed without knowledge of treatment group.

### 2.4 Imaging and analysis

The spinal cords stained with immunofluorescence were imaged on a Zeiss confocal using 20X 0.8 NA Plan Apo (image size 425.1μm x 425.1μm) and 40X 1.2 NA C Apo (image size 212.5μm x 212.5μm) objectives. Images were acquired as 16 bit and 1024x1024 dimensions. Images were taken from dorsal, ventral, and two lateral locations in axial sections of the spinal cord. CD4, CD11b and Iba1 images were quantified using thresholding method in MetaMorph (7.10 version) software. The data was represented as % stained area (Chaudhary *et al.* 2011). ASPA labeled cells were imaged using a z stack and the images were converted to a single maximum intensity projection images using Zen Black software. ASPA positive cells were counted and represented at per mm^2^.

Plastic embedded semi-thin spinal cord sections were imaged using Zeiss Imager 2 microscope at 10X 0.45 NA Plan Apochromat, 20X 0.8 NA Plan Apochromat and 63X 1.4 NA Plan Apochromat (198x159 μm) using black and white Axiocam 506 mono camera. Images at 20X were stitched together to get the entire spinal cord montage using Zen software. The images were 300-400MB so they could be digitally zoomed to look at axons providing enough details for analysis. The entire ventrolateral area of each section was analyzed for damage (areas with absence of axons, decompacted myelin with separation of lamellae and degenerating axons that contained electron dense axoplasm and absence of cytoplasmic details, Chaudhary et al. 2011) using MetaMorph software. The damaged area was totaled and represented as % damaged area. The ultra-thin spinal cords sections on nickel grids were stained using 5% uranyl acetate and Reynold’s lead citrate, dried and imaged using FEI Techani T12 electron microscope. EM images were acquired from lateral spinal cord. Eight images (18.5 x 14.5 microns, 2900X) were collected from both sides of each spinal cord. Two out of the 8 images were randomly chosen for analysis. The total axons counted were classified into normal myelinated, degenerated and unmyelinated. The axons were plotted across all groups. All imaging analyses were done blinded to the treatment condition.

### 2.5 Oligodendrocyte progenitor cell (OPC) culture

OPCs were purified from postnatal rat brains according to the procedure described by Dugas and Emery 2013. Briefly, OPCs were purified by positive selection using O4 immunopanning after removal of astrocytes and OL using Ran2 and anti-GalC, respectively. Purity of the cultures was ~95% as verified by O4 staining. Purified OPCs were expanded in poly-d-lysine coated flasks for 4-5 days in SATO serum-free media containing Platelet-Derived Growth Factor-AA (PDGF-AA, 10 ng/ml, PeproTech), neurotrophin-3 (NT-3, 1 ng/ml, PeproTech, Rocky Hill, NJ), and ciliary neurotrophic factor (CNTF,10 ng/ml, PeproTech). For differentiation and survival assays, OPCs were passaged and plated onto poly-d-lysine coated glass coverslips in 24 well plates at 50,000 cells per well in SATO serum-free media containing NT-3 (1 ng/ml, PeproTech), and CNTF (10 ng/ml, PeproTech). OPC differentiation was induced by withdrawal of PDGF for 48 hours in the absence of T3 and cells were then treated with either 0.1% DMSO (vehicle control), 50nM T3 or Sob for a further 24 hours. For assessment of differentiation, cells were fixed for 10 minutes in 4% paraformaldehyde and stained overnight with anti- 2’, 3’-cyclic nucleotide 3’- phosphodiesterase (CNP, Millipore MAB326) and proteolipid protein (PLP, AA3 rat hybridoma, kind gift of Richard Reynolds) in 0.3% Triton X-100 and 10% normal goat serum in PBS. The coverslips were washed and stained with Alexa Fluor 488- conjugated anti-mouse and 555- conjugated anti-rat secondary antibodies for two hours at room temperature. Coverslips were mounted with Fluoromount-G with DAPI nuclear stain (ThermoFisher). Images were taken from six independent coverslips on a Zeiss Axio Imager M2 using a 20X objective and the proportion of viable cells positive for PLP and CNP determined by a scorer blind to the experimental condition. Dead cells were identified by condensed and/or fragmented DAPI-stained nuclei. For survival assays, cells were incubated for 10 minutes in calcein AM and ethidium homodimer using the Live/Dead viability assay (ThermoFisher) per manufacturer’s instructions. Live (green) and dead (red) cells were imaged using an IncuCyte ZOOM analysis system (Essen Biosciences Inc, Ann Arbor, MI). Four images per well, 3 wells/condition were manually scored for the number of live and dead cells by a blinded observer.

### 2.6 Statistical analysis

All analyses were done in a blinded manner and the mice were randomly assigned to vehicle or treatment groups. Statistical differences were compared between vehicle and treatment groups using Mann Whitney U tests using Graphpad, Prism (San Diego, CA). A p value of 0.05 was considered to be significant. All p values and number of mice used in each group are indicated in the figure legends. Vehicle mice (50% DMSO and 8mM NaOH) were pooled for analysis. For disease scores, statistical differences were assessed using a one-way ANOVA with a Tukey’s multiple comparison test for significance. For cell culture experiments, statistical differences were assessed using a one-way ANOVA with a Bonferroni’s multiple comparison test for significance.

## 3. Results

### 3.1 T3, Sob and Sob-AM2 reduced Total EAE scores and damage to the spinal cord

The suppression effects of T3, Sob and Sob-AM2 were assessed in a widely used murine EAE model in C57BL/6 mice. EAE was induced in mice using MOG 35-55 peptide where the average onset of disease was at day 15 post-immunization in mice receiving vehicle. Compared with vehicle, T3, Sob, and Sob-AM2 significantly reduced EAE disease severity (Fig 1A,B, Table 1). The mean total EAE scores ± SE were vehicle 31.2±3.6, T3 19.1 ± 4.0 (p<0.05), Sob 10.8 ± 3.1 (p <0.0009), and Sob-AM2 5mg/kg 4.3 ± 1.9 (p <0.0009). One-way ANOVA was performed for total EAE score and the p value for all groups was <0.0001. Further analysis using Tukey’s multiple comparisons were done (Sob, T3 and SobAM2 groups compared to vehicle were significant p value 0.0005, 0.04, and <0.0001 respectively, Table 1). While T3, Sob and Sob-AM2 consistently suppressed clinical EAE, initiating treatment after onset of signs of clinical EAE did not alter clinical course or pathology (data not shown).

**Fig 1.**
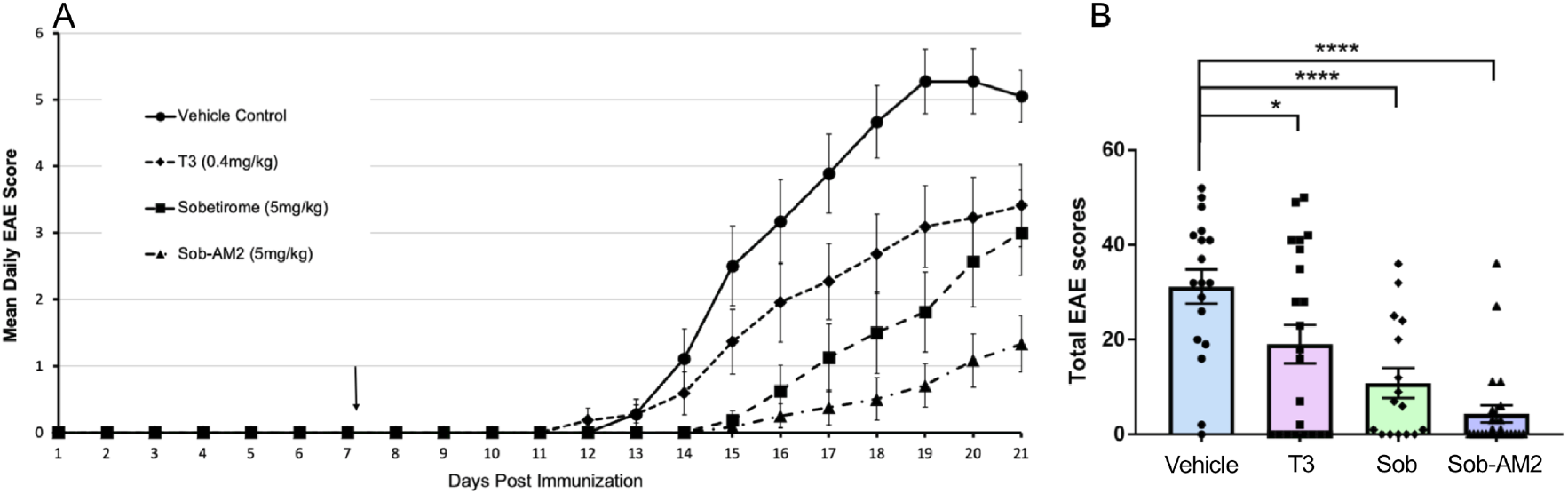
Lower disease scores and damage in T3, Sob and Sob-AM2 groups. Daily EAE scores (A) and Total EAE scores (B) in various groups were plotted. All values represented as averages ± SE. p value <0.0009**** and <0.05*. The graphs include data from vehicle n=18, T3 n=22, Sob n=16, and Sob-AM2 5mg/kg n=24. Data is a composite of three independently experiments performed following the same protocol.

### 3.2 T3, Sob and Sob-AM2 reduced spinal cord damage and preserved myelinated axons

We evaluated the lumbar spinal cords for axonal/myelin damage using toluidine blue staining. Ventro-lateral total area was measured and the areas with absent axons, degenerated axons and decompacted myelin were marked as damaged areas (Fig 2A-D). Damaged area was expressed as a percent of ventro-lateral white matter. Mice receiving vehicle had the highest percent damage at 9.7±1.6 and Sob-AM2 5mg/kg had the least amount of damage at 2.2 ± 1.1 (p <0.0009, Fig 2A’-D’, E). Percent damage ± SE for T3 was 6.4 ± 1.8 (p<0.05); and for Sob was 5.2 ± 2.0 (p<0.02). All statistical comparisons were done with vehicle group. EM of spinal cord sections demonstrated significantly more normal appearing myelinated axons in mice receiving T3, Sob and Sob-AM2 compared with vehicle treated mice (Fig 3A-F). T3, Sob and Sob-AM2 also showed a significant decrease in degenerated axons (Fig 3G). No difference was detected in the number of completely unmyelinated axons and unmyelinated axons were similar in number to that of naïve mice, suggesting that primary demyelination had not occurred as previously described for this EAE model (Jones *et al*. 2008). The healthy myelinated axons had normal morphologic appearance. Mean percent normal appearing myelinated axons + SE: vehicle 42.8± 4.3, T3 62.4 ± 3.9 (p<0.005), Sob 62.8 ± 5.2 (p<0.02), Sob-AM2 68.0± 4.0 (p <0.0009) and naïve 74.8 ± 3.0. Mean percent degenerated axons + SE: vehicle 39.6 ± 3.7, T3 24.3 ± 3.5 (p<0.005), Sob 22.2 ± 4.0 (p<0.005), Sob-AM2 5mg/kg 20.0 ± 3.6 (p <0.0009) and naïve 7.3 ± 0.9. Thus T3, Sob and Sob-AM2 significantly decreased myelin and axon degeneration compared with vehicle.

**Fig 2.**
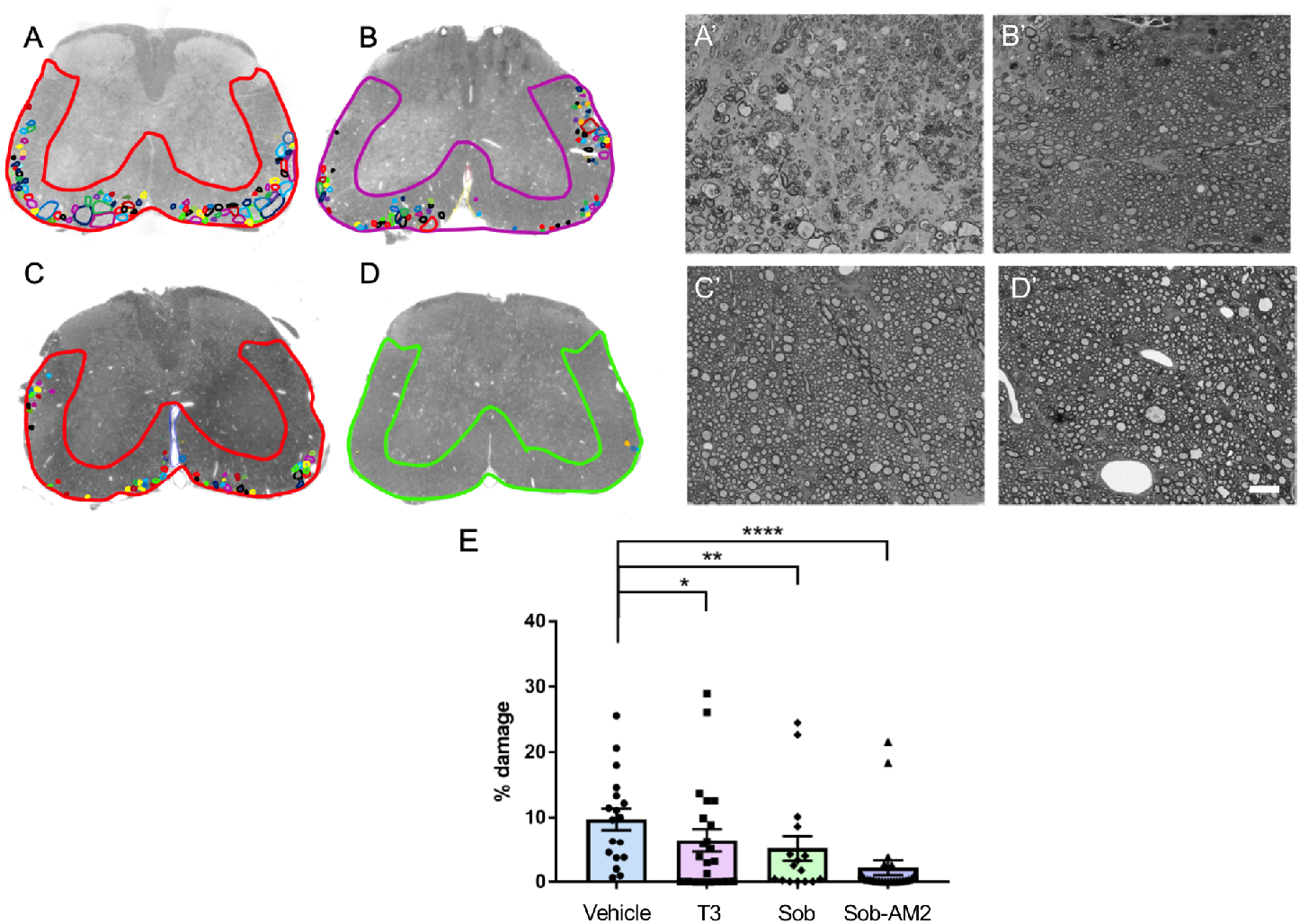
Reduction of damage in ventrolateral spinal cord. Representative damage in A,A’ Vehicle; B,B’ T3; C,C’ Sob; D,D’ Sob-AM2. Ventrolateral area and small areas of damage are marked to get percent damage. The graphs include data from vehicle n=18, T3 n=22, Sob n=16, and Sob-AM2 5mg/kg n=24. F. plot of damage in all the groups. p value <0.0009****, <0.02** and <0.05*.

**Fig 3.**
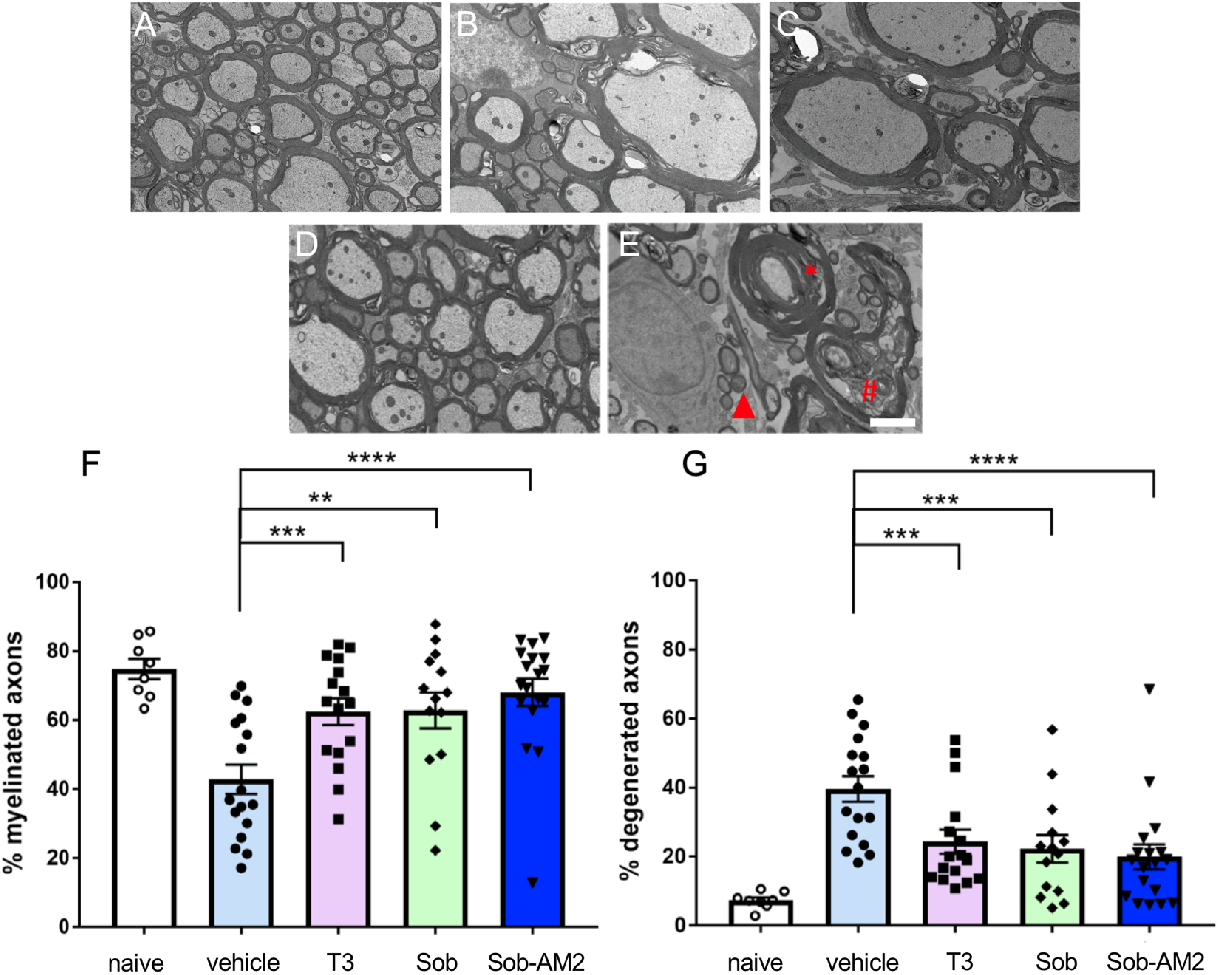
Higher number of healthy axons and less number of degenerated axons were detected in treated groups. A. Naïve; B. T3; C. Sob; D. Sob-AM2; E. vehicle Red asterisk shows decompacted myelin, # axon with debris and triangle shows degenerated axon. F. quantitation of myelinated axons G. quantitation of degenerated axons. p value <0.0009****, < 0.005***, and <0.02**. The graphs include data from vehicle n=17, T3 n=16, Sob n=14 Sob-AM2 n=18 and naïve n=8. Scale bar 2 μm.

### 3.3 Thyromimetics effects on CD4, CD11b, and Iba1 staining

To address whether the tissue protective effects of T3 and the thyromimetics were related to reduction in inflammatory cells, we studied the effects on the populations of CD4 positive T cells, and CD11b and Iba1 positive microglia/macrophages in T3, Sob and Sob-AM2 treated mice as compared to vehicle. This analysis was performed in the lumbar spinal cords at the end of the EAE time course (day-21). CD4-positive T cells were affected by T3 and Sob-AM2 treatment, which significantly lowered the T cell population relative to vehicle control whereas Sob had CD4-positive cell numbers that did not differ significantly from vehicle control (Fig 4A-D). Percent CD4 staining ± SE values were vehicle 1.1 ± 0.2, T3 0.6 ± 0.1 (p=0.05), Sob 0.7 ± 0.2 (n.s.), and Sob-AM2 0.4 ± 0.1 (p <0.0009) (Fig 4E).

**Fig 4.**
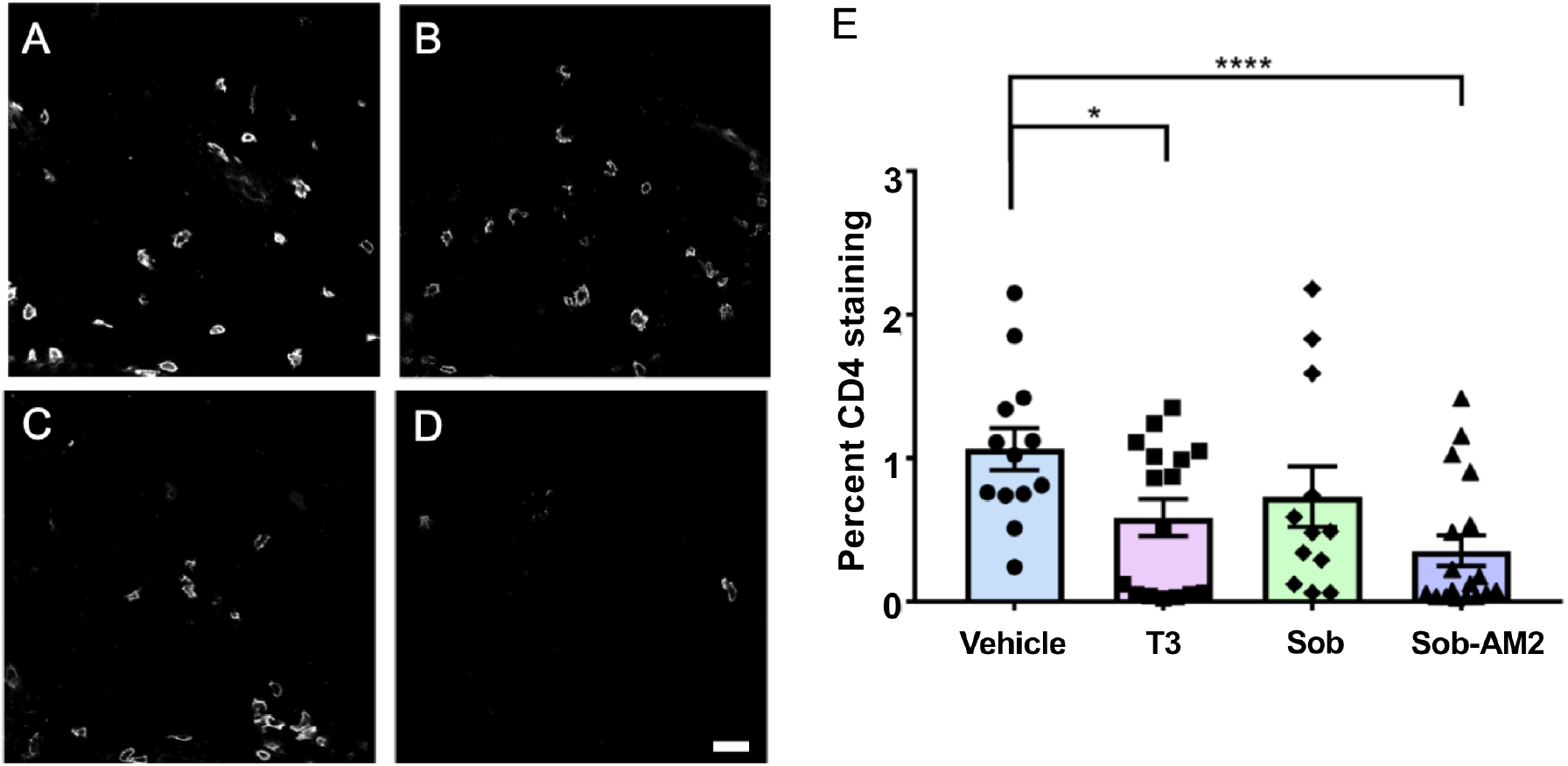
Reduction in CD4 positive T cells. Representative CD4 images from T3, Sob and Sob-AM2 groups show decrease in levels of CD4 cells. A. Vehicle; B. T3; C. Sob; D. Sob-AM2 (Scale bar 20 μm); E. quantitation of CD4 staining. p value <0.0009**** and <0.05*. The graphs include data from vehicle n=13, T3 n=16, Sob n=12, and Sob-AM2 n=18.

Both CD11b+ and Iba1+ microglia/macrophage population were significantly reduced by Sob and Sob-AM2 treatment. T3 treatment also significantly decreased CD11b staining but did not significantly affect Iba1 staining (Fig 5A-L). Percent CD11b staining ± SE values were the following: vehicle 9.6 ± 1.2; T3 5.5 ± 1.2 (p<0.05); Sob 4.1 ± 1.2 (p<0.005); Sob-AM2 2.6 ± 0.8 (p <0.0009, Fig 5M). Percent Iba1 staining ± SE values were the following: vehicle 8.7 ±1.2; T3 6.2 ± 1.5; Sob 4.7 ± 1.2 (p<0.05); and Sob-AM2 2.3 ± 0.5 (p<0.0009, Fig 5N).

**Fig 5.**
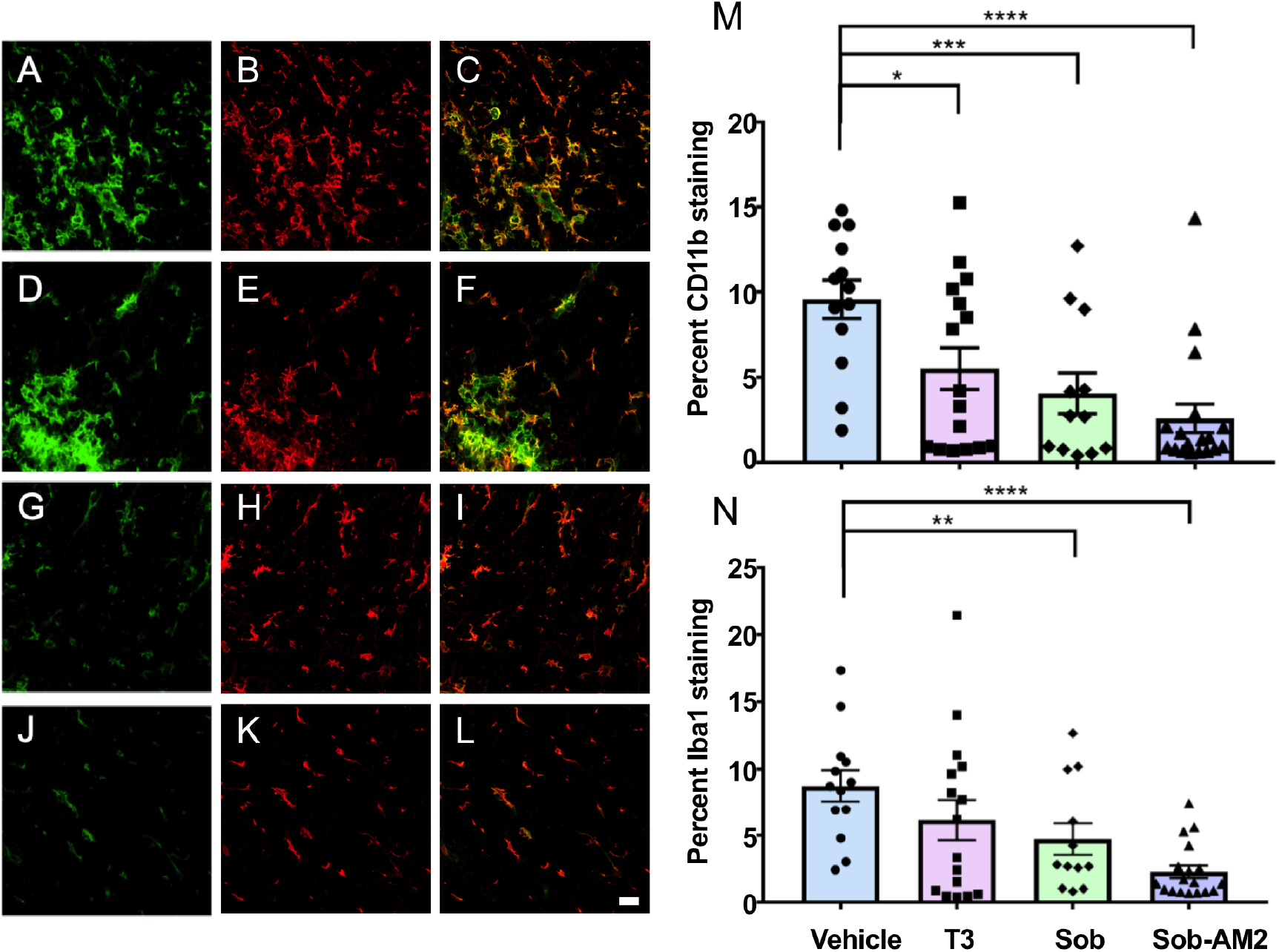
Reduction in CD11b and Iba 1 positive microglia and macrophages after treatment with T3, Sob and Sob-AM2. Representative CD11b and Iba1 double stained spinal cord images. Green CD11b, red Iba1 and overlay of both staining is shown. A,B,C vehicle; D,E,F T3; G,H,I Sob; J,K,L Sob-AM2 (Scale bar 20 μm). M. CD11b quantitation; and N. Iba1 quantitation. p value <0.0009****, <0.005***, <0.02** and <0.05*. The graphs include data from vehicle n=13, T3 n=16, Sob n=12, and Sob-AM2 n=18.

### 3.4 CNS selective thyromimetics decrease loss of mature ASPA labeled oligodendrocytes

In order to assess whether treatment with T3, Sob or the Sob-AM2 prodrug reduced loss of oligodendrocytes in EAE, we assessed the density of mature oligodendrocytes, using ASPA as a marker at the Day 21 time-point. Relative to naïve mice, vehicle treated EAE mice showed a significant reduction in the density of ASPA+ oligodendrocytes, consistent with the expected myelin degeneration and oligodendrocyte loss (1137 ± 66 cells/mm^2^ in naïve vs. 800 ± 60 cells/mm^2^ in vehicle treated EAE mice, p<0.02, Fig 6). In contrast, drug treated groups did not show a reduction in ASPA+ oligodendrocyte densities relative to the naïve controls (p>0.05 for all comparisons). Both the T3 (1005 ± 64 cells/mm^2^) and Sob-AM2 (1098 ± 61 cells/mm^2^) treated animals showed a significant increase in OL densities relative to vehicle treated EAE mice (p<0.05 and p<0.005, respectively), restoring them to naïve control levels. Sob had higher number of ASPA+ cells as compared with vehicle but the difference did not reach significance (p= 0.1).

**Fig 6.**
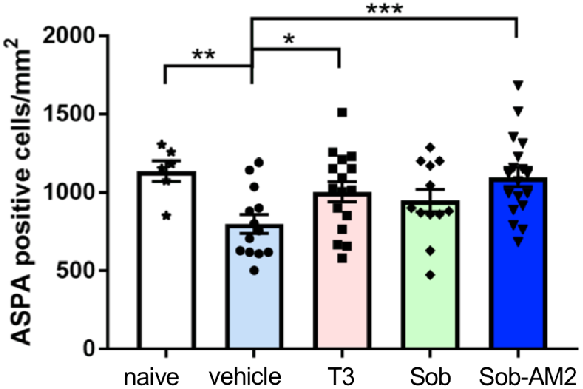
T3 and Sob-AM2 treatment preserved ASPA+ oligodendrocyte numbers. Although there was an increase in the OL in Sob treated groups these numbers did not reach significance. A graph of ASPA counts/mm^2^ in the different groups. p value <0.005***, <0.02**, and <0.05*. The graphs include data from vehicle n=13, T3 n=16, Sob n=12, Sob-AM2 n=18 and naïve n=6. Scale bar 20 μm.

### 3.5 Thyromimetics increase the survival of mature OL in cell culture

In addition to thyroid hormone’s role in promoting OPC differentiation (Billon *et al.* 2002; Schoonover *et al.* 2004; Barres *et al.* 1994), it has also been shown to promote the survival of oligodendrocytes (Jones *et al.* 2003). The relative preservation of ASPA+ oligodendrocytes along with normal appearing myelin in the T3, Sob and Sob-AM2 treated EAE mice suggested that these agents may be partially acting through an oligodendrocyte protective mechanism. We therefore assessed the ability of T3 and Sob to enhance oligodendrocyte survival in culture. Rat OPCs were differentiated for 48 hours in the absence of PDGF, by which time the majority of cells are multipolar CNP+ OLs. The OLs were then exposed to vehicle (0.1% DMSO), 50nM T3 or 50nM Sob for a further 24 hours and their viability assessed (Fig 7A). It has previously been noted that immature OLs are highly susceptible to apoptosis, with ~50% of developing oligodendrocytes undergoing apoptosis during development (Barres *et al*. 1992; Trapp *et al*. 1997). Cultured oligodendrocytes display a similar vulnerability to apoptosis. Consistent with previously published observations (Jones *et al.* 2003), T3 exposure increased viability of the oligodendrocytes from 59.7 ± 2.4% in the vehicle condition to 80.9 ± 1.7% (p<0.005). Sob promoted oligodendrocyte survival equally well (84.3 ± 1.5%, p<0.005, Fig 7B). Staining of the cultures with an early (CNP) and late (PLP) maker for differentiation confirmed that the majority of cells in all conditions were mature PLP+ oligodendrocytes (Fig. 7C). When only viable cells were assessed the proportion of cells positive for PLP was significantly increased by both T3 and Sob (46.9 ± 5.1 in vehicle, 67.0 ± 2.0 in T3 and 72.5 ± 3.5 in Sob, p<0.005 and <0.02 for T3 and Sob, respectively). Although this difference may in part reflect promotion of differentiation by T3 and Sob, the majority of dead cells (identified by pyknotic or condensed nuclear and fragmented processes) were PLP+ (Fig. 7C). This suggests that the difference in the proportion of viable cells that were PLP+ may be accounted for by the selective increase of survival of the maturing oligodendrocytes. The ability of T3 and Sob to promote survival of oligodendrocytes *in vitro* may result from increasing the resistance of maturing oligodendrocytes to apoptosis to which they are vulnerable (Barres *et al.* 1992; Sun *et al.* 2018). Together, these results indicate that like T3, Sob can promote the viability of OLs *in vitro.*

**Fig 7.**
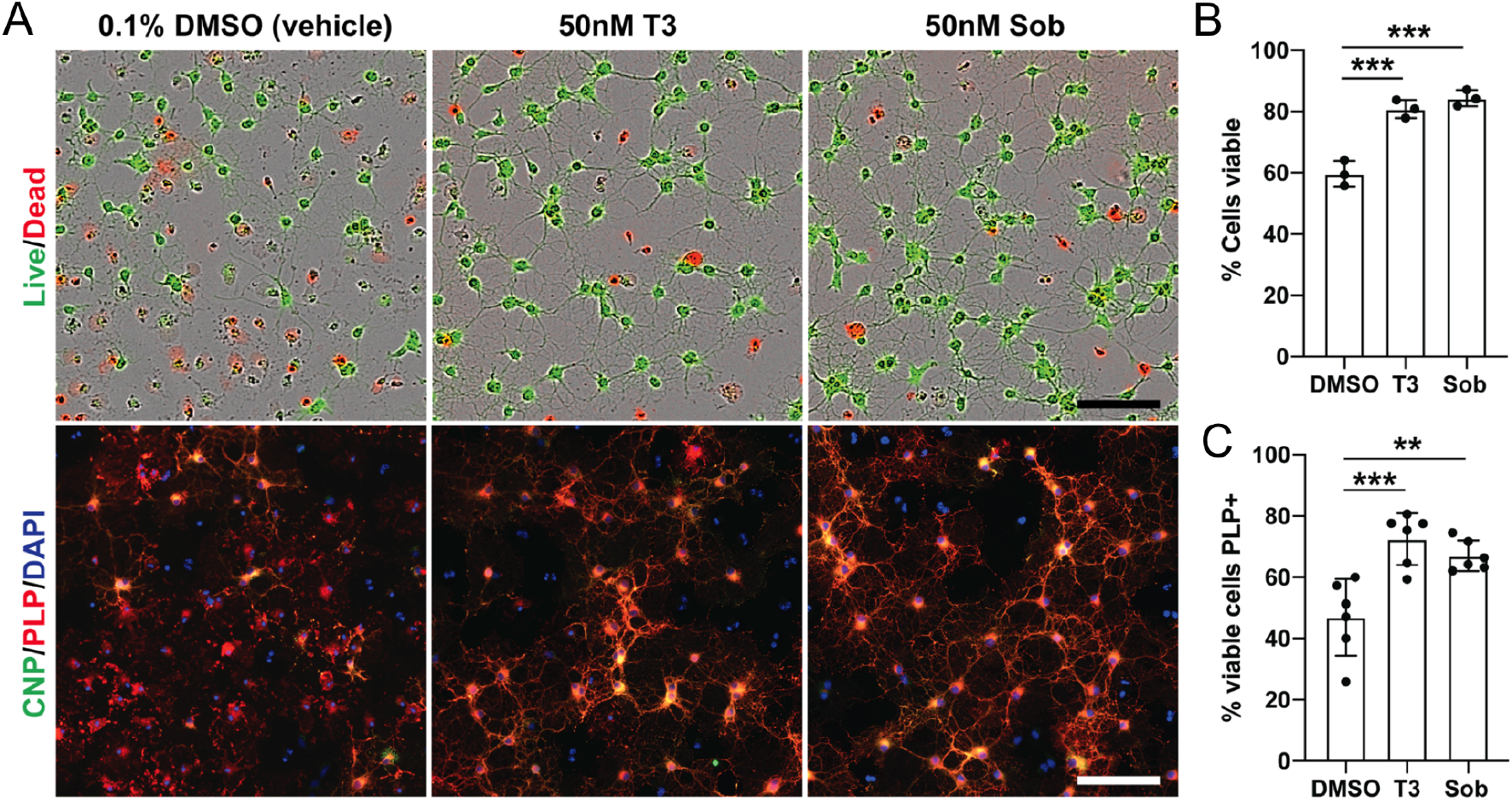
Oligodendrocyte survival *in vitro* after exposure to T3 and Sob. T3 and Sob protect OL against cell death in culture. Rat OPCs were differentiated for 48 hours by withdrawal of PDGF before being cultured for a further 24 hours in vehicle (0.1% DMSO), 50nM T3 or 50nM Sob. A. Representative images from each condition stained to assess viability (Live/Dead) or differentiation (anti-CNP and PLP). B. Quantification of the proportion of viable cells in each condition, determined by the Calcein AM/Ethidium homodimer Live/Dead assay. C. Quantification of the proportion of viable cells in each condition that are mature (PLP+) OLs. p value <0.005*** and <0.02**. Scale bars = 100μm.

## 4. Discussion

This study confirms previous studies on the therapeutic benefit of thyroid hormone in EAE, and for the first time demonstrates the therapeutic effects in EAE of a TRβ selective thyromimetic, Sob, and its CNS penetrating prodrug Sob-AM2. Importantly, we demonstrate the protection of myelin, oligodendrocytes and axons in EAE by T3 and the thyromimetics. Of interest is that we have recently demonstrated the ability of Sob and Sob-AM2 to stimulate remyelination in non-EAE murine models of CNS demyelination (Hartley *et al.* 2019). The current study thus broadens the treatment potential for Sob and Sob-AM2 in MS, suggesting that thyromimetics may be neuroprotective as well as being able to stimulate remyelination.

The mechanisms by which T3, Sob, and Sob-AM2 reduce myelin degeneration, oligodendrocyte loss and axonal degeneration are likely to be complex (Figs 2,3). All cell types express thyroid hormone receptors and therefore have the potential for T3 and the thyromimetics to influence their function. The effects of T3 on oligodendrocyte maturation are well known and we and others have shown that Sob can stimulate differentiation of OPC into oligodendrocytes (Baxi *et al.* 2014). We demonstrate here that Sob, like T3, can promote survival of oligodendrocytes *in vitro* (Fig 7) and demonstrate a protective effect in EAE mice exposed to T3 and Sob (Fig 6). Protection of oligodendrocytes by T3 and Sob would be expected to reduce loss of myelin and also indirectly protect axons. Thus, we propose that protection of oligodendrocytes may be one important mechanism to explain the positive effects of T3 and Sob in EAE, although effects on other CNS cell types, such as astrocytes, need to be considered. It is important to note that, astrocytes play an important role in TH metabolism *in vivo*, influencing the local levels of T3 through deiodination of T4 via type 2 deiodinase (Morte and Bernal 2014). In addition, TH signaling can enhance astrocyte glutamate uptake (Mendes-de-Aguiar et al. 2008), providing a protective effect against glutamate-mediated toxicity. We demonstrate here that Sob, like T3, can promote survival of oligodendrocytes *in vitro* and propose that a similar direct pro-survival effect may occur in EAE mice exposed to T3 or Sob. Nevertheless, since the *in vitro* studies used purified OPCs, the beneficial effects of T3 and Sob *in vivo* may also involve indirect effects mediated by astrocytes or other cell types. The relative contributions of different cell types (such as oligodendrocytes, astrocytes and immune cells) to the beneficial effects of T3 and Sob will be important to establish in future work, perhaps through conditional ablation of TH receptors.

We also identified effects on inflammatory cells in EAE mice receiving T3, Sob and Sob-AM2 (Fig 4 and 5). The effects on CD4+ T cells were not consistent. Compared with vehicle, T3 had a modest but significant lowering of CD4+ cells and Sob did not have a significant lowering of CD4+ cells whereas Sob-AM2 reduced CD4+ cells. Despite these differences in effects on CD4+ cells, all three treatments decreased EAE severity, reduced tissue injury and inhibited myelin and axonal degeneration. As such, we believe the beneficial effects of T3, Sob and Sob-AM2 cannot be explained solely by an effect on CD4+ cells. All three agents however caused reductions in CD11b+ microglia/macrophages and Sob and Sob-AM2 reduced expression of an activation marker, Iba1, on microglia/macrophages. Microglia and macrophages are effector cells in EAE and actively involved in phagocytosis of myelin. TH is known to play a role in innate immunity and TH receptors are expressed by both macrophages and microglia (van der Spek *et al.* 2017). So the effects of T3, Sob and Sob-AM2 on CD11b+ cells might be related to reduction in the number and activation state of these cells. The effect of Sob and Sob-AM2 but not T3 on Iba1 expression suggest that TRß selective thyromimetics may inhibit microglial activation more effectively than T3. It is also possible that the reduction in CD11b+ cells could be secondary to the reduction in myelin degeneration. Myelin fragments are known to activate microglia and monocytes (Williams *et al.* 1994; Kopper and Gensel 2018) and so the changes in CD11b+ cells in mice receiving T3, Sob and Sob-AM2 could result from their inhibition of myelin fragmentation. The importance of anti-inflammatory effects to the therapeutic benefit of T3, Sob and Sob-AM2 in EAE is uncertain and warrant further investigation, particularly the apparent effects of Sob and Sob-AM2 on microglial activation.

Despite the uncertainty of how T3, Sob and Sob-AM2 modulate EAE, the effects are impressive and are at least in part consistent with a neuroprotective effect with particular inhibition of myelin and axon degeneration and oligodendrocyte loss. Of the three treatments, Sob-AM2 had the most profound effects compared with T3 and Sob. Sob-AM2 has increased penetrance of the blood brain barrier compared with Sob and, once gaining access to the CNS, fatty acid amide hydrolase cleaves the amide side-chain from the prodrug to produce high levels of Sob in the CNS (Meinig et al. 2017) Sob-AM2 produces low levels of Sob in the circulation so it has an even lower potential of producing systemic thyrotoxic side-effects than the parent compound. Further investigation is needed to elucidate the mechanisms of the neuroprotective effect of Sob and its prodrug, Sob-AM2. Given its ability to produce high concentrations of Sob within the CNS, its low potential to cause systemic thyrotoxicosis and its neuroprotective effects and ability to stimulate remyelination, Sob-AM2 and related compounds warrant consideration as a treatment for MS.

## Declarations

### Ethical Approval and Consent to participate

The VA Portland Health Care System (VAPORHCS) IACUC committee approved all experiments on mice.

### Consent for publication

Not applicable.

### Availability of supporting data

All data generated or analyzed during this study are included in this published article

### Funding

National Multiple Sclerosis Society

RG-5199A4 and RG-1607-25053 to DB, RG-5106A1/1 to BE NIH DK52798 to TSS

NIH P30 NS061800

Race to Erase MS to DB

OHSU Laura Fund for Innovation in Multiple Sclerosis to DB and TSS

### Authors’ contributions

DB, TSS and BE conceived the experiments. PC, GM designed the experiments. GM and EP provided animal care, disease induction and monitoring, performed perfusions and analyses. PC and EC conducted IHC and EM processing and analyses. BE cultured OPC. AB analyzed the OPC data. PC wrote the manuscript with suggestions from DB, TSS, BE and GM.

## Acknowledgements

This research was supported by the National Multiple Sclerosis Society grants RG 5199A4 and RG-1607-25053 to DB, RG 5106A1/1 to BE), the Race to Erase MS to DB, the OHSU Laura Fund for Innovation in Multiple Sclerosis to DB and TSS and NIH DK52798 to TSS. We would like to thank Advanced Light Microscopy Core and Electron Microscopy Core supported by the NIH, P30 NS061800. We would also like to thank the VA Portland Health Care System for providing animal husbandry, equipment support and laboratory space.

## List of abbreviations

EAE: experimental autoimmune encephalomyelitis
ip: intraperitoneal
MOG: myelin oligodendrocyte glycoprotein
OPC: oligodendrocyte precursor cell
Sob: sobetirome
TH: Thyroid hormone
T3: TH, 3,5,3’-triiodothyronine

